# The First Publicly Available Annotated Genome for Cannabis plants

**DOI:** 10.1101/786186

**Authors:** Conor Jenkins, Ben Orsburn

## Abstract

Recently we have seen a relaxation of the historic restrictions on the use and subsequent research on the *Cannabis* plants in North America. We have recently described a pipeline for the creation of annotated protein databases using a combination of genomic and proteomic data and the application of this method toward the analysis of the proteomes of *Cannabis* plants. In parallel with our work, we approached the National Center for Biotechnology Information (NCBI) to align and annotate publicly deposited genomic files for these plants. In September of 2019, this project was completed. The result, NCBI Cannabis sativa Annotation Release 100 is now live and publicly available. The annotated genome allows, for the first time, the use of classical genetics and proteomics tools for the interrogation of these plants. Furthermore, the presence of an annotated genome within the NCBI Genome browser now permits any researcher with a web browser to manually examine or BLAST genetic sequences, vastly reducing the resources required to create primers to amplify genes from the plants or microbial contaminants that may affect them. We describe this new resource and some points of obvious value to the scientific community as well as the integration into the *Cannabis* Proteome Draft Map Project.

**Significance Statement:** Until recently laws in North America have restricted nearly all research on *Cannabis* plants. Until recent research from our lab, only a few hundred genes and proteins from the plant had been annotated for putative function. The construction of a publicly available annotated genome for this plant allows, for the first time, the use of traditional genomic and proteomic tools for the investigation of these plants. We evaluate herein the first fully annotated publicly available genome for *Cannabis* plants and the integration of this resource into www.CannabisDraftmap.org

## Introduction

*Cannabis* plants, generally regarded as *Cannabis sativa* and *Cannabis indica* have been grown for centuries for industrial, medicinal and recreational purposes. The near complete global restrictions on these plants in the early 20^th^ century has severely limited any research into these organisms. The passage of the Farm Act of 2019 in the United States has allowed, nation-wide access to *Cannabis* plants that possess a total concentration of THC of less than 0.3% weight. Coupled with decriminalization and legalization of all *Cannabis* plants in Canada and an increasing number of states, the interest in *Cannabis* spans multiple areas of medical and industrial science. We have recently described the first attempt at global proteomic characterization of these plants. A major challenge in the Cannabis Draft Map Project was the lack of a publicly available and fully annotated genome for these plants. In June of 2019, we approached the National Center for Biotechnology Information (NCBI) to request they utilize publicly deposited genomes from these plants to create the first fully annotated reference for the *Cannabis* genome. The NCBI accepted this request and the result, NCBI Cannabis sativa Annotation Release 100, was announced as complete and publicly available on September 9^th^, 2019. This resource allows, for the first time, interrogation of annotated *Cannabis* genetics by any researcher through any common genomics or proteomics tools. Furthermore, by hosting this resource on the interactive NCBI Genome Browser, any interested research with a web browser may search and evaluate this valuable resource. We have utilized these newly available materials to both improve and refine our current draft map data of these plants and here compare the completed annotations to our in-house generated resources.

## Results and Discussion

The generation of the EggNOG annotated protein FASTA and shotgun proteomics data was described in detail in our recent investigation.^1^ The resources generated by the NCBI follow a well-established pipeline that has been extensively described in these publicly available resources.^2–4^ Table 1 provides the web addresses for all data that will be described in this text.

**Table 1.** Direct links to all data described in this study

| Resource description | Web Address |
| --- | --- |
| *Cannabis* Proteomics Data | https://www.cannabisdraftmap.org |
| EggNOG Annotated FASTA | https://www.cannabisdraftmap.org/resources |
| **NCBI Cannabis sativa Annotation Release 100** |  |
| Annotated Cannabis Genomes | ftp://ftp.ncbi.nlm.nih.gov/genomes/all/GCF/900/626/175/GCF_900626175.1_cs10 |
| Annotation Statistics | https://www.ncbi.nlm.nih.gov/genome/annotation_euk/Cannabis_sativa/100/ |
| Genome Browser Direct Link | https://www.ncbi.nlm.nih.gov/genome/gdv/browser/?acc=GCF_900626175.1&context=genome |

**Table 2.**
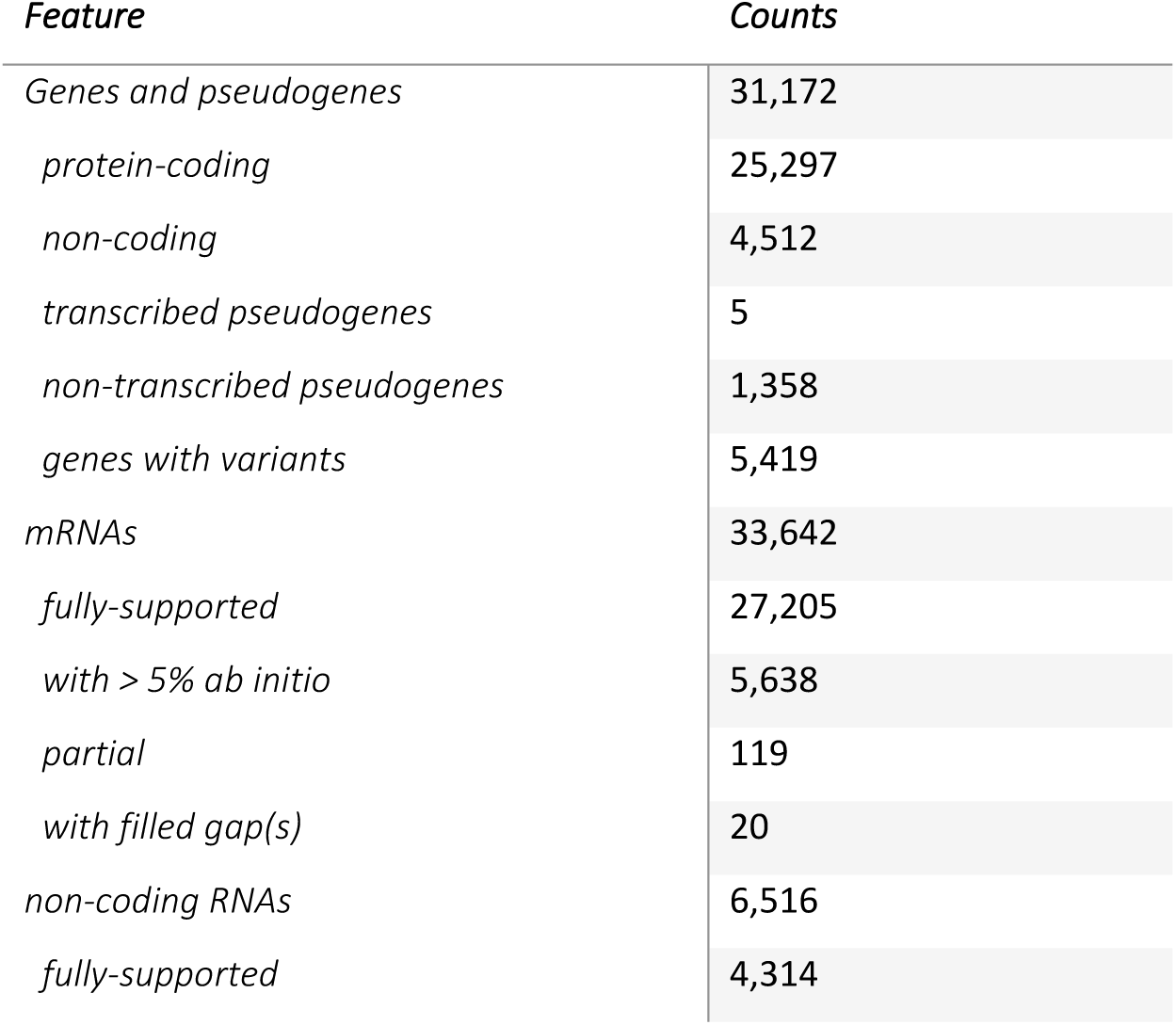
Feature counts from this genome assembly as described.

While many groups are currently working on *Cannabis* genomics using varying technologies, including both long and short read sequencing, relatively few groups have made the raw data publicly available.^5^ Even when this data may be downloaded, the evaluation of unprocessed data is often beyond the capabilities of many researchers, as most code requires access to high performance computer clusters for analysis. The release of this annotation by the NCBI is a great advance for researchers without the expertise or computational resources to perform these analyses on their own.

One major advantage is the presence of the annotated genome deposition with direct accessibility through the NCBI Genome Browser. This tool allows the direct examination of genomics information to anyone within a web browser and internet access, as all data processing is performed on the NCBI internal servers.^6^ Figure 1 is an image taken from this resource and a high resolution PDF image is available as Supplemental Figure 3. One immediate value of the presence of the complete annotated genome within the Genome Browser is the access to the NCBI Primer BLAST tool. Primer BLAST allows, for the first time, the rapid and design and quality control of primers designed to amplify specific Cannabis sativa genes. Furthermore, this tool allows for primers designed for the detection of pathogenic microorganisms to be referenced for possible matching sequences within the *Cannabis* genome. Nearly any lab can now design specific high-quality primers for *Cannabis* genetics and microbial contamination work.^7^

**Figure 1.**
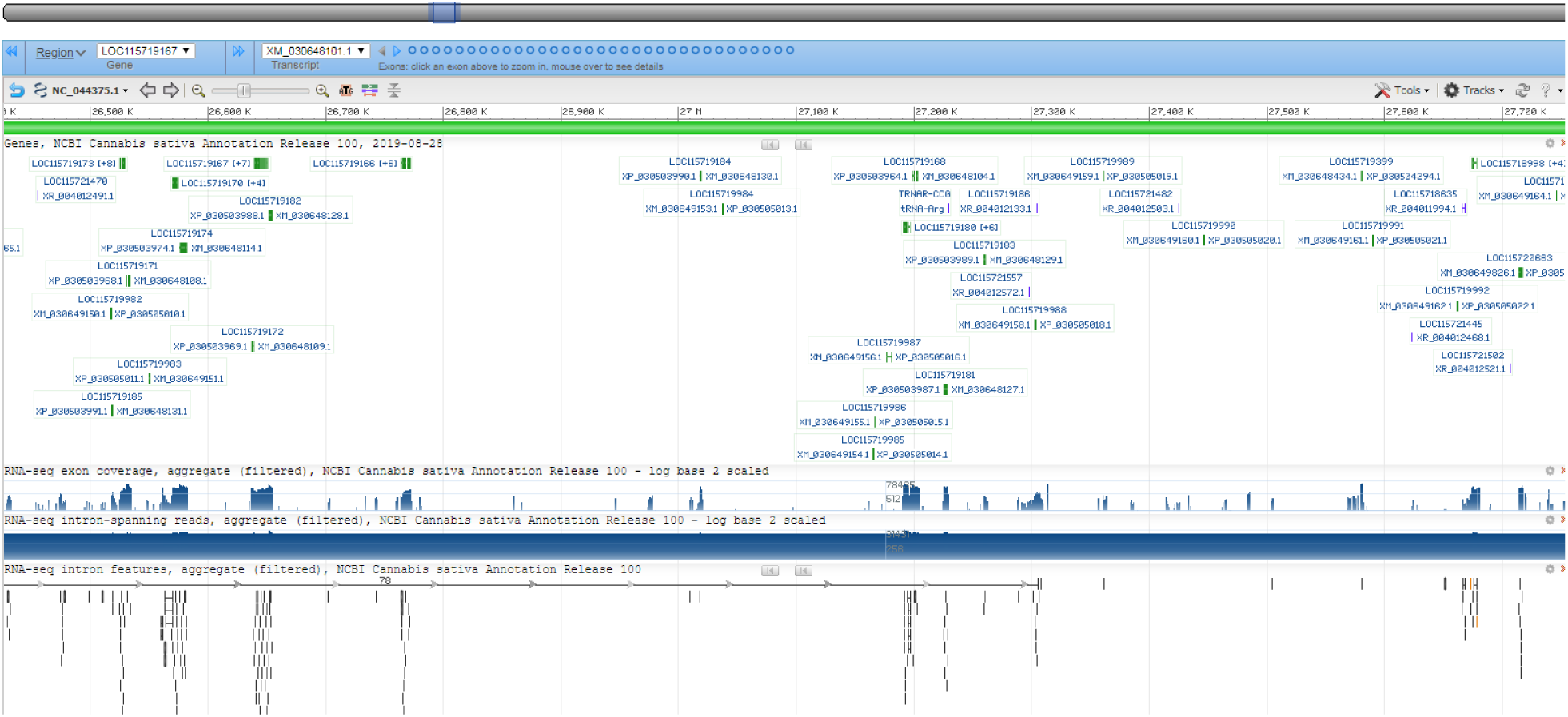
An image taken from the Cannabis sativa Annotation Release 100 visualized in the publicly available NCBI Genome Browser.

Proteomics data analysis relies, almost entirely, on the comparison of tandem mass spectra (MS/MS) to annotated genomics data. While proteomic data analysis may be performed using FASTA data directly obtained from sequencers, few supercomputers or Cloud based proteomic search engines currently exist to make these analyses truly feasible due to their expense and lack of widespread access. Nearly all processing for proteomics is performed on desktop computers using reduced, annotated, often manually reviewed, and often targeted theoretical protein FASTA databases (reviewed in Bolt ^8,9^). The release of the NCBI annotations essentially permits, for the first time, both shotgun and top-down analysis of proteomics data from *Cannabis* plants.

Using a high-performance computer designed for and dedicated to the task, we completed a proteogenomic analysis of Cannabis plant material, resulting in both a theoretical proteins FASTA database and peptide/protein identifications. When utilizing the direct genomics data, individual search iterations required over 100 hours of searching per search iteration. The use of the EggNOG or NCBI FASTAs allow a full search of all data currently available for the project to be performed in less than 10 hours using the same hardware (data not shown).

**Table 3.** Features in NCBI Cannabis sativa Annotation Release 100

| *Feature* | *Count* | *Mean length*<br>*(bp)* | *Median*<br>*length (bp)* | *Min length*<br>*(bp)* | *Max length*<br>*(bp)* |
| --- | --- | --- | --- | --- | --- |
| *Genes* | 29,809 | 3,477 | 2,343 | 63 | 976,063 |
| *All transcripts* | 40,158 | 1,651 | 1,431 | 63 | 17,828 |
| *mRNA* | 33,642 | 1,791 | 1,566 | 114 | 17,396 |
| *misc_RNA* | 1,229 | 2,034 | 1,779 | 146 | 14,917 |
| *tRNA* | 491 | 74 | 73 | 71 | 91 |
| *lncRNA* | 3,085 | 1,079 | 605 | 73 | 17,828 |
| *snoRNA* | 1,595 | 106 | 107 | 63 | 243 |
| *snRNA* | 99 | 142 | 129 | 64 | 196 |
| *rRNA* | 17 | 903 | 155 | 115 | 3,394 |
| *Single-exon*<br>*transcripts* | 4,528 | 1,206 | 993 | 114 | 5,625 |
| *coding transcripts* | 4,528 | 1,206 | 993 | 114 | 5,625 |
| *CDSs* | 33,677 | 1,373 | 1,140 | 114 | 16,644 |
| *Exons* | 155,404 | 315 | 167 | 1 | 9,202 |
| *in coding transcripts* | 145,964 | 317 | 168 | 1 | 7,968 |
| *in non-coding*<br>*transcripts* | 13,151 | 262 | 138 | 2 | 9,202 |
| *Introns* | 123,569 | 535 | 156 | 30 | 970,509 |
| *in coding transcripts* | 117,315 | 518 | 153 | 30 | 970,509 |
| *in non-coding*<br>*transcripts* | 9,792 | 694 | 202 | 32 | 81,832 |

To directly compare the protein FASTA files from generated in our previous study to the new NCBI annotations we performed 3 identical searches with the only alteration being the protein FASTA database(s) utilized. An overview of these results is shown in Table 4. When comparing results when protein uniqueness is determined by a single peptide to define a protein group, the EggNOG FASTA database comes out clearly ahead of both the NCBI database, as well as when both resources are searched in tandem. This apparent advantage is reduced markedly when a minimum of 2 unique peptides are required to define a high confidence protein.

**Table 4.** Proteomic analysis comparing the results of searching the complete *Cannabis* proteome draft versus the two annotated protein FASTA files.

| *FASTA Database* | *PSMs* | *Protein groups* | *Protein groups with >1 unique peptide* |
| --- | --- | --- | --- |
| *EggNOG v1.1* | 233,921 | 14112 | 7503 |
| *NCBI Can Sat Release 100* | 203,224 | 8001 | 6151 |
| *Both Databases* | 256,203 | 9729 | 9735 |

When both databases are used in tandem, more group identifications are found to originate from the NCBI database, as shown in Supplemental Figure 1. When the molecular weights of the proteins detected by each individual FASTA database are plotted in kilodalton (kDa), the distributions suggest that the EggNOG FASTA matches result from consistently smaller theoretical protein sequences, as demonstrated in Figure 2 and Supplemental Figure 2. Taken together, these results suggest that many of the assignments made by the initial analysis of the *Cannabis* proteome may be contributed to incorrect assignments of start or stop sites. However, over 2,809 new protein assignments can still be confidently assigned to the EggNOG FASTA when used in tandem with the NCBI database, suggesting that individual genetic variation or alternative open reading frames may still play an important role in the differences between these results.

As an evaluation of peptide spectral match (PSM) confidence, the individual database search data was binned by cross correlation score (XCorr), as shown in Figure 2. XCorr is a common and well-characterized metric for the confidence of peptide matches. Surprisingly, no statistical difference was found between the XCorr distribution of the two sets, with the EggNOG having an arithmetic mean of 2.73 and SD of 0.93 and the NCBI output of 2.60 and SD 0.97. A distribution of the XCorr of each individual search, as well as of the PSMs uniquely attributed to the EggNOG FASTA when both databases are used in tandem is shown in Figure 2. Further manual review will be necessary to determine the true size of the Cannabis proteome. This is not surprising considering the number of proteins in the human proteome is still under debate after decades of continuous study.^10^

**Figure 2.**
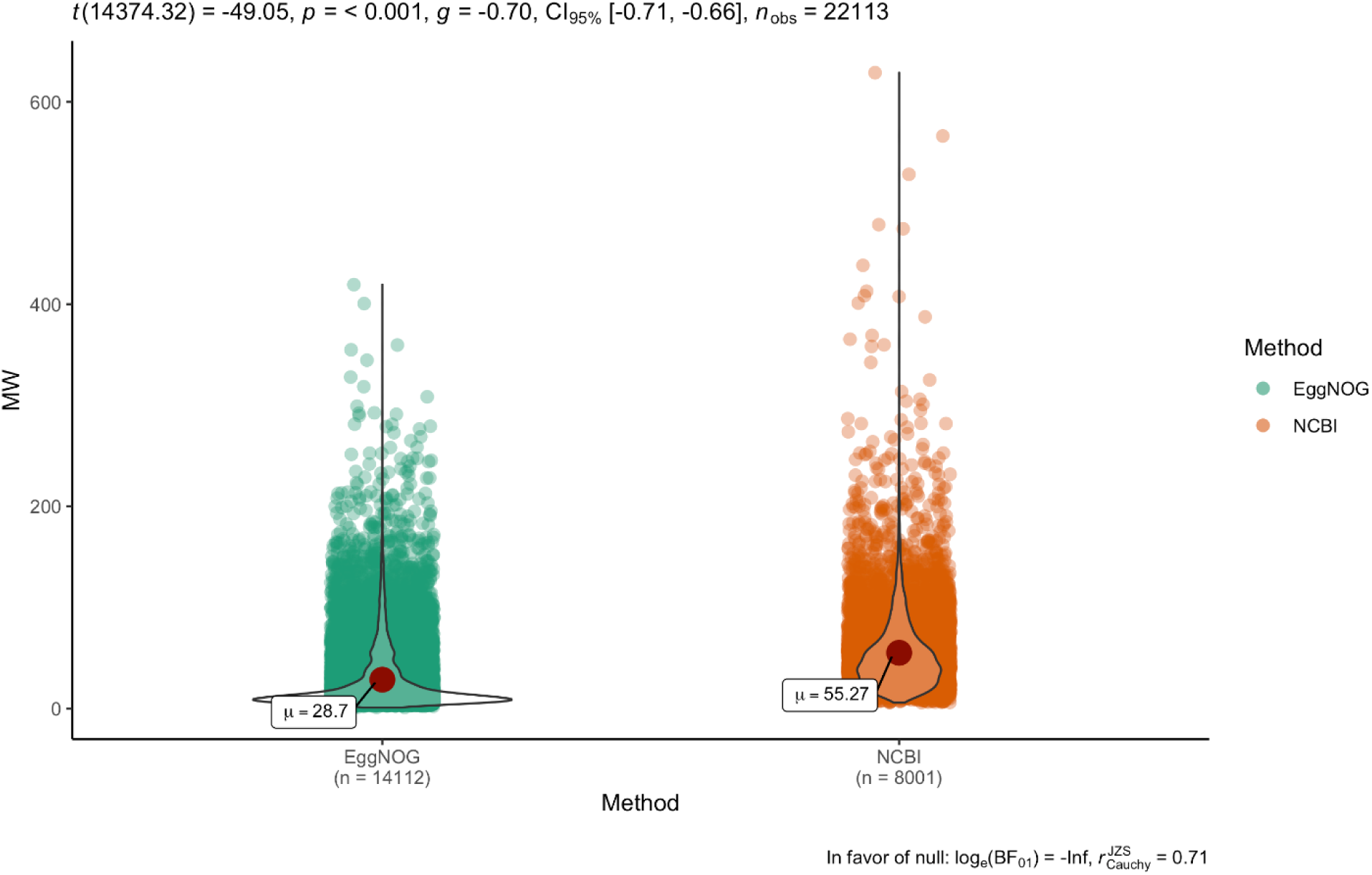
A combined plot of MW vs numbers of proteins as violin plot. Median and outliers are marked as described.

**Figure 2.**
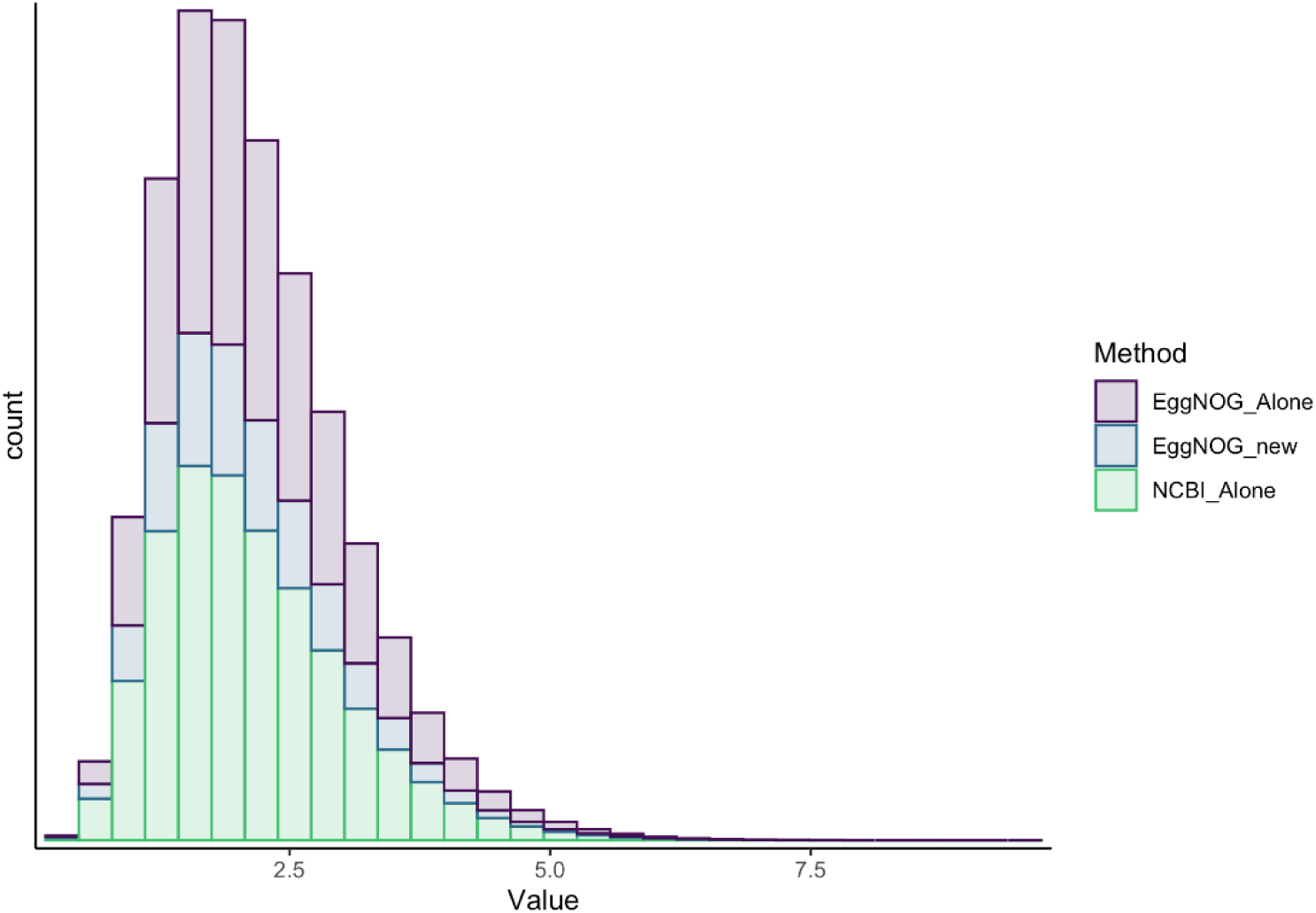
A distribution of the cross-correlations (XCorr) for the peptides attributed to each database shown

## Conclusions

We have described herein some of the details and value of the first publicly available and fully annotated *Cannabis* genome, NCBI Cannabis sativa Annotation Release 100. This resource permits, for the first time, the use of all common and web-based tools for genetics and proteomic analysis to be utilized for studies of Cannabis plants. We have also demonstrated how this resource is allowing the improvement and refinement of our work profiling the *Cannabis* proteome. All resources described here are live and openly available.

## Acknowledgements

The creation of the NCBI Cannabis sativa Annotation Release 100 was performed, in it’s entirety by the staff scientists of the NCBI Eukaryotic Genome Assembly Pipeline. We would like to thank Dr. Miranda Darby of Hood College School of Bioinformatics for conversations helpful to the construction of this work.

## Supplemental Methods

### Comparison of annotated protein databases

All high resolution files from the Cannabis Proteome Draft were downloaded in MGF format from www.CannabisDraftMap.org and searching in Proteome Discoverer 2.2 using a vendor default pipeline consisting of SequestHT and Percolator. For all searches, a 10ppm MS1 tolerance and 0.02 Da MS/MS tolerance were used along with the static modification of iodoacetamide carbamidomethylation of cysteines and the dynamic modification of oxidation on methionine. Three searches were performed, with each search containing the cRAP database of common contaminants. The first search used the EggNOG FASTA alone, the second was performed with NCBI Release 100 and the third search with all 3 databases in tandem. All plots were generated in R Studio using ggPlot2. All code is uploaded here as Supplemental data.

## Supplemental Figures

**Supplemental Figure 1.**
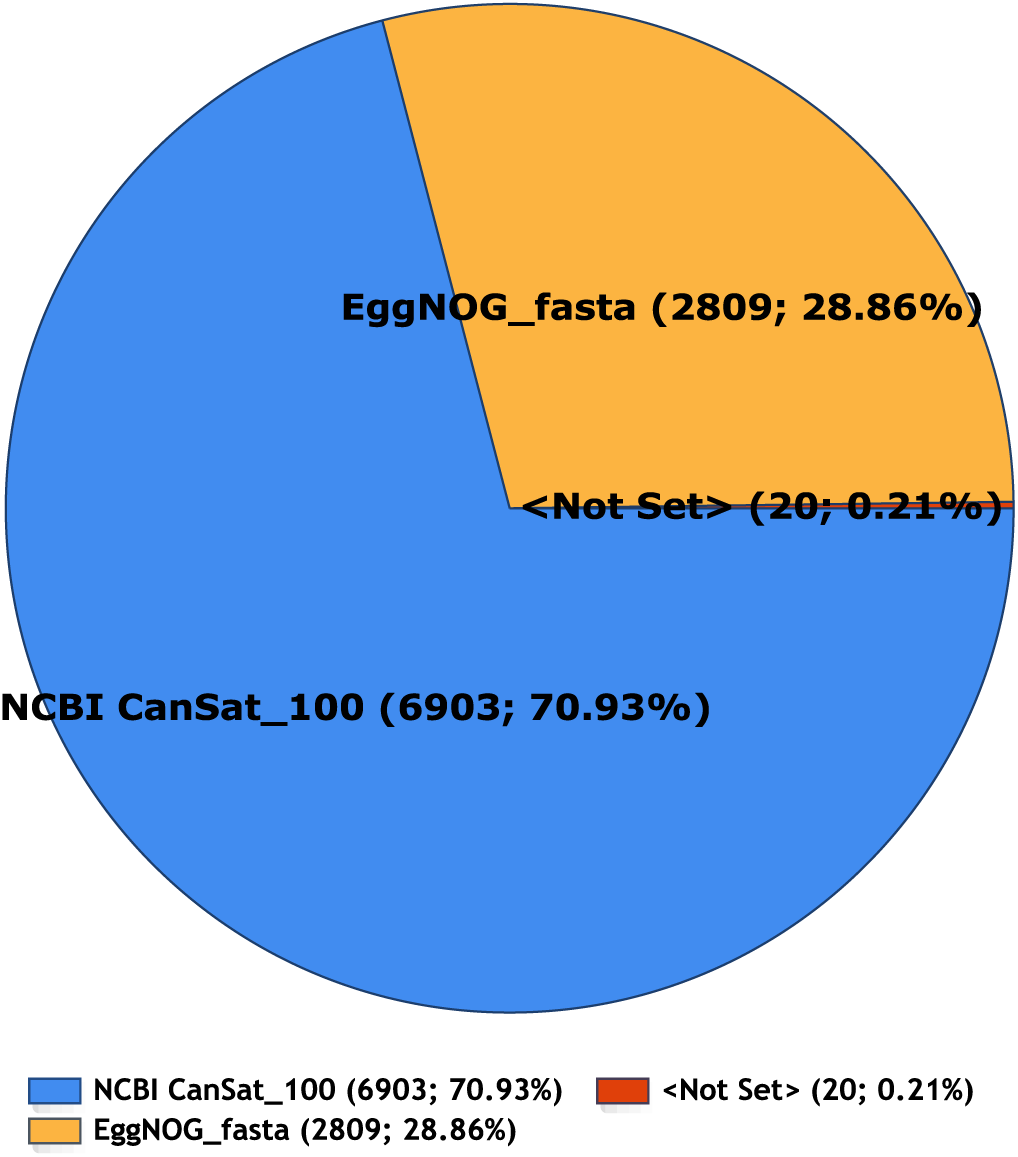
A pie chart representing the number of annotations assigned to each theoretical protein database when both resources described in this study are utilized in tandem.

**Supplemental Figure 2.**
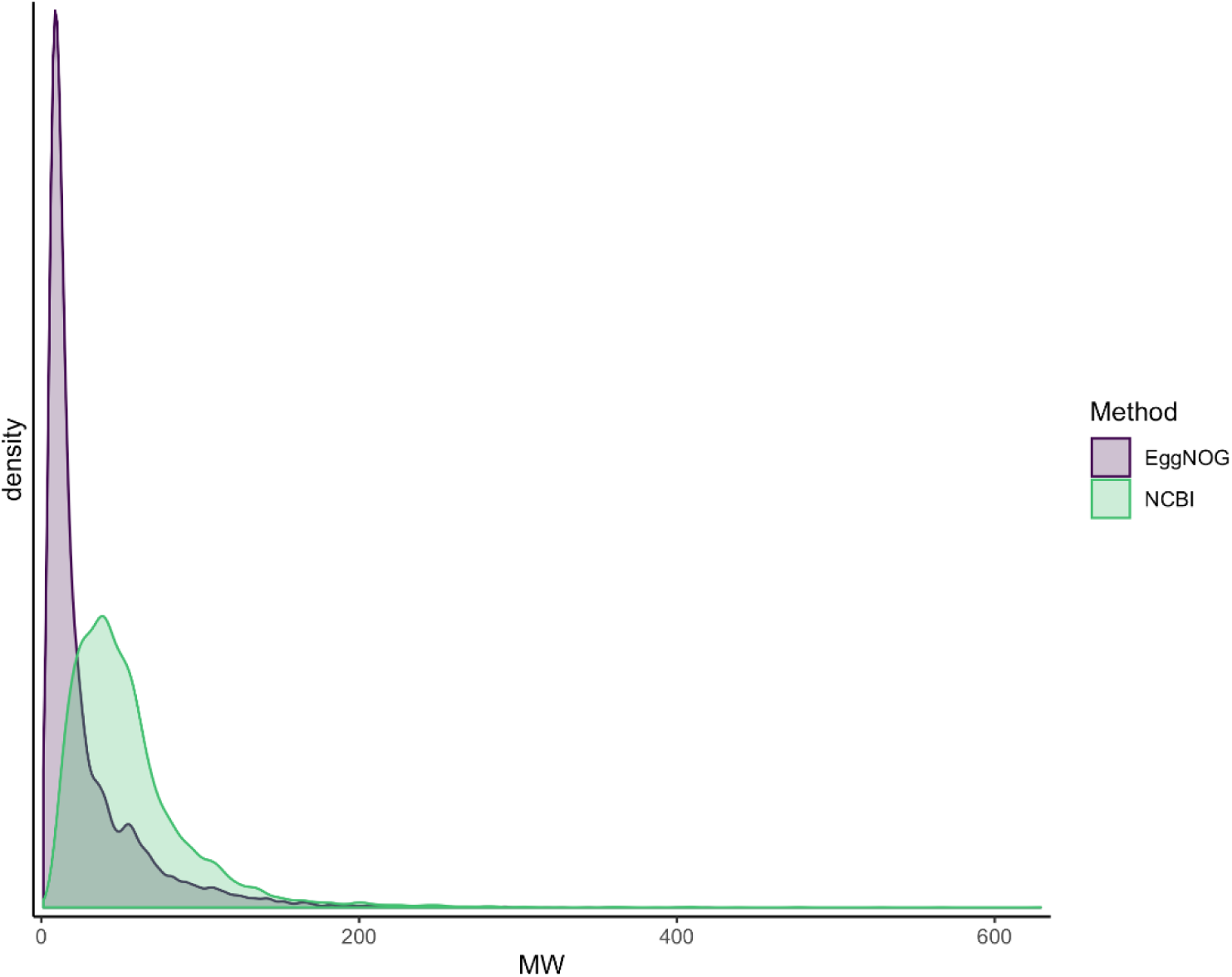
A density plot of protein molecular weights assigned to the individual protein searches versus each FASTA database when searched separately.

