## Supplementary figures and images for "The First Publicly Available Annotated Genome for Cannabis plants"

### Supplemental Figure 3

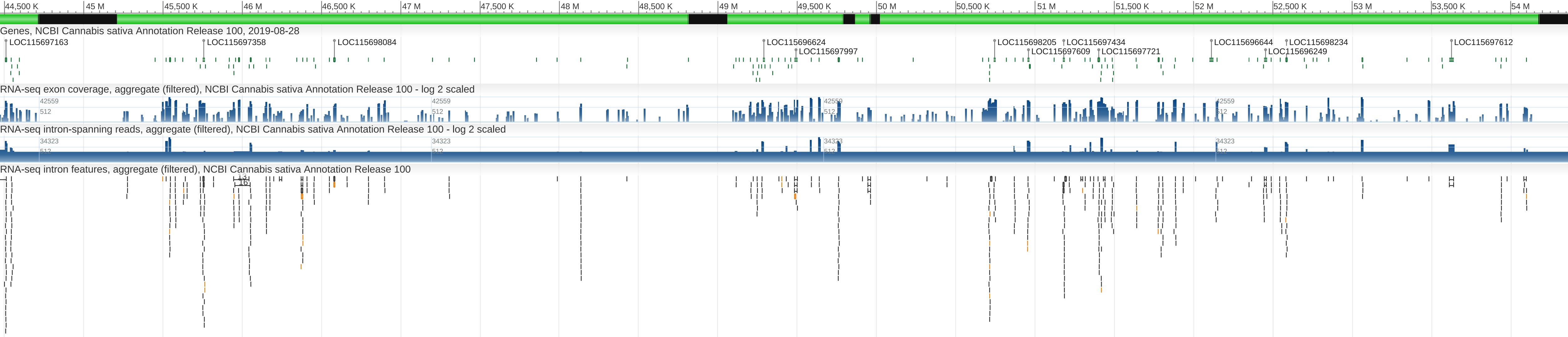
